# Standardized and calibrated light stimuli via head-mounted displays for investigating the non-visual effects of light

**DOI:** 10.1101/2025.01.15.633125

**Authors:** Maydel Fernandez-Alonso, Manuel Spitschan

**Affiliations:** Translational Sensory & Circadian Neuroscience, Max Planck Institute for Biological Cybernetics, Tübingen, Germany; Chronobiology & Health, TUM School of Medicine and Health, Technical University of Munich, Munich, Germany; TUM Institute for Advanced Study (TUM-IAS), Technical University of Munich, Germany

## Abstract

Light influences human physiology profoundly, affecting the circadian clock and suppressing the endogenous hormone melatonin. Experimental studies often employ either homogenous full-field stimulation, or overhead illumination, which are hard to standardize across studies and laboratories. Here, we present a novel technique to examine non-visual responses to light using virtual-reality (VR) head-mounted displays (HMDs) for delivering standardized and calibrated light stimuli to observers in a reproducible and controlled manner. We find that VR HMDs are well-suited for delivering standardized stimuli defined in luminance and across time, with excellent properties up to 10 Hz. We examine melatonin suppression to continuous luminance-defined light stimuli in a sample of healthy participants (n=32, mean±SD age: 27.2±5.6), and find robust melatonin suppression in 24 out of 32 participants (75% of the sample). Our findings demonstrate that VR HMDs are well-suited for studying the mechanisms underlying human non-visual photoreception in a reproducible and standardized fashion.

## Introduction

### The non-visual effects of light

Light plays a fundamental role in allowing us to visually perceive the world around us, but it also exerts a profound influence on other physiological and behavioural processes in humans such as circadian entrainment, regulation of hormone secretion, and modulation of alertness and mood (Blume et al., 2019; Brown et al., 2022). These non-visual effects are predominantly mediated by a specialized subset of retinal cells containing the photopigment melanopsin, the intrinsically-photosensitive retinal ganglion cells (ipRGCs) (Berson et al., 2002; Dacey et al., 2005; Do, 2019; Do et al., 2009; Hattar et al., 2002; Lucas et al., 2014; Spitschan, 2019). Over the past two decades, numerous studies have demonstrated that light exposure at specific intensities and wavelengths can acutely suppress the hormone melatonin, affect alertness levels, and shift the timing of the circadian clock (Brainard et al., 2001a; Brown, 2020; Brown et al., 2022; Cajochen et al., 2000; Cajochen, 2007; Khalsa et al., 2003; Lewy et al., 1980; Phillips et al., 2019; Thapan et al., 2001; Zeitzer et al., 2000). As such, precisely controlled light interventions have become a key tool for researchers investigating a range of outcomes, from basic circadian biology to clinical applications in sleep and mood disorders.

### Current methods for light stimulation and limitations

In circadian and sleep research, the most common approaches to deliver light stimuli include overhead room illumination, light boxes or ganzfeld dome stimulators (Spitschan et al., 2023), often equipped with multi-channel LED light sources that allow researchers to vary the intensity levels and spectral compositions of the illumination (Barrionuevo et al., 2022; Brainard et al., 2001b; Thapan et al., 2001). More recently, some studies have also incorporated digital displays as a light source in evening studies to deliver a controlled light intervention (Blume et al., 2024; Schöllhorn et al., 2023).

Despite the widespread adoption of these techniques, they carry several limitations. When using overhead room illumination, participants’ head movements, eye movements and changes in posture can cause variability in the amount of light actually reaching the retina (Spitschan, 2021; Thibos et al., 2018), leading to uncontrolled differences in exposure that may confound experimental outcomes. Second, in more controlled setups, participants are required to remain in carefully controlled conditions while limiting considerably their posture and head movements, reducing ecological validity and the comfort of the participant. The need for carefully controlled laboratory conditions also precludes the possibility to conduct at-home studies. Third, the nature of these setups often imposes limitations in cost, laboratory space or both, having the consequence that data collection is limited to one participant per evening, significantly slowing the rate at which studies can be completed.

### Head-mounted displays for controlled stimulus presentation

The past decade has witnessed a rapid development of virtual and augmented reality technologies. From their early development in the mid-2010s, head-mounted displays (HMDs) have experienced exponential growth, with a very wide variety of affordable and more capable devices available today. While most HMDs have been oriented towards the consumer market, researchers have been quick to take advantage of these new technologies as powerful instruments for conducting behavioural research with humans. Virtual reality (VR) creates an immersive three-dimensional environment that can be fully controlled by the experimenter. Improvements in display hardware, such as better resolution and greater field of view, ensure a more precise stimulus presentation and improve immersion. Furthermore, advancements in rendering technologies and the increased accessibility of software platforms to create realistic virtual environments, has made developing experiments in VR increasingly common (Creem-Regehr et al., 2023).

Beyond recreating real environments, HMDs can also help solve some of the problems with traditional experimental setups, particularly in the field of visual perception. For a typical screen setup, it is crucial to maintain a fixed distance and viewing angle between the observer and the screen, which is often achieved by highly restricting the participant’s posture and head movements with the use of chin rests or similar. Furthermore, it is often important to control that no visual stimulation is presented concurrently, which creates the need for completely dark environments where one can ensure that no other stimulus will appear in the observer’s field of view. As a result, such setups are often only feasible to implement in a laboratory, making them impractical for other applications outside this setting. Since HMDs are strapped to the observer’s face, the distance between the eyes and the display remains constant, and their immersive nature ensures that researchers have full control of the stimuli that appear within the field of view of the observer. These advantages have led to HMDs being used for psychophysical experiments (Creem-Regehr et al., 2022; Cwierz et al., 2021; De Gelder et al., 2018; Lepori et al., 2023), clinical studies (Mees et al., 2020; Neugebauer et al., 2021; Tatiyosyan et al., 2020; Wiley et al., 2022), and also for applications outside the lab, such as at-home vision testing (Chia et al., 2023; Hu et al., 2023; Tsapakis et al., 2018).

However, one untapped field where the use of HMDs could be advantageous is in light exposure protocols that investigate the non-image-forming effects of light. By delivering a standardized, calibrated light stimulus that encompasses most of the participant’s field of view and remains constant regardless of changes in posture or head movements, HMDs open new possibilities for exploring the mechanisms underlying the non-visual effects of light. One specific application is the study of binocular integration, as HMDs enable precise, independent control of the light delivered to each eye. This capability, combined with the potential to present complex spatially or temporally modulated stimuli, goes beyond what is currently achievable with overhead room illumination or other existing methods.

While VR technology has been used in sleep and circadian research (Goldsworthy et al., 2023), its applications have mostly been for therapeutic or behavioural interventions, rather than as a tool for delivering controlled light stimuli. Previous studies have shown that HMDs displays can emit light with spectral characteristics capable of influencing melatonin secretion (Kim et al., 2018; Wu et al., 2019). However, these studies focused on theoretical modelling or device characterization through light measurements rather than testing physiological effects in real participants. Our study builds on this by directly assessing melatonin suppression in participants exposed to light via a VR headset.

To our knowledge, this is the first study to experimentally demonstrate the physiological effects of light exposure delivered through a VR HMD in humans. The integration of precise display calibration with biological outcome measures provides a foundation for using VR HMDs as a controlled light delivery tool in circadian research.

### Integrated eye tracking technology

In more recent times, advancements in HMDs have included integrated eye-tracking capabilities, which make it possible to precisely monitor gaze direction and pupil size in real-time (Adhanom et al., 2023). This feature is particularly relevant to studies on the non-visual effects of light, as pupil size is directly modulated by the ipRGCs, and at the same time it directly determines retinal irradiance, influencing the magnitude of physiological responses (Gaddy et al., 1993; Higuchi et al., 2008; Schöllhorn et al., 2024). Moreover, tracking gaze direction can help researchers verify compliance (i.e., confirming that participants have their eyes open and are looking at a stimulus for the intended duration). With eye tracking becoming more accessible, the integration of sophisticated calibration and measurement tools within VR HMDs paves the way for a new generation of controlled, individualized light exposure studies that can be performed inside and outside traditional laboratory settings.

### Current study

The purpose of the present study is to develop and validate a novel method for delivering standardized, calibrated light stimuli using VR HMDs. We implement temporal and spectral calibration procedures to characterise the displays in use, and implement an evening light exposure intervention to test their feasibility for studying acute melatonin suppression. Ultimately, this approach has the potential to broaden the methodological possibilities within the field of chronobiology, enabling new avenues of research into the non-visual effects of light.

## Materials and Methods

### Apparatus: Hardware and software setup

Stimuli were presented using a virtual reality (VR) head-mounted display (HMD), the HTC Vive Pro Eye (HTC, Taiwan). This device integrates a high-resolution binocular display with built-in Tobii eye-tracking technology capable of real-time pupillometry and eye movement measurements. Stimulus control and presentation were managed using custom code developed in Python, using the licensed program Vizard 7 from WorldViz (Santa Barbara, CA, USA). The VR headset allowed for precise control of light stimuli, ensuring calibrated and reproducible light exposure across participants.

The HTC Vive Pro Eye has dual 3.5-inch Organic Light-Emitting Diode (OLED) displays, one for each eye. The technical specifications given by the manufacturer are:

- Resolution: 1440 x 1600 pixels per eye (2880 x 1600 combined)
- Refresh rate: 90 Hz
- Field of view: 110 degrees

### Calibration of head-mounted display

#### Luminance calibration and gamma correction

A luminance calibration of the HMD was performed to ensure a linear light output. Measurements were taken using a Jeti spectraval 1511-HiRes spectroradiometer (Jena Technische Instrument, Jena, Germany) to measure spectral radiance and luminance for each screen primary (RGB) and for white at various input values (from 0 to 1 in steps of 0.1). These measurements were repeated for both displays (i.e. left and right eye). The measurements were processed and fitted using a power function, such that:

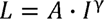

where *L* represents the measured luminance, *I* is the input, *A* is a scaling factor, and γ is the gamma. The inverse of this exponent was then used to model and correct the non-linear behaviour of the display, such that:

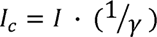

Where *I* is the desired input (e.g., 0.5 for 50% intensity), γ is the estimated gamma value, and *I_c_* is the corrected input to use. We verified the effectiveness of the gamma correction by re-measuring the luminance of the display after correction and confirming a linear luminance output.

#### Illuminance and α-opic equivalent measurements

In addition to spectral radiance, we measured the spectral irradiance at the eye position to derive the corneal illuminance and α-opic equivalents. The measurements were taken using a Jeti spectraval 1501 spectroradiometer (Jena Technische Instrument, Jena, Germany). To ensure consistent positioning across the different virtual reality headsets tested and the reproducibility of measurements, the irradiance measurements were taken using a custom-designed 3D-printed adapter tailored for the HTC Vive Pro Eye (see **Figure 1**, left). This adapter aligns the spectroradiometer with the headset’s eye relief point, minimizing angular variation and ensuring repeatable measurements across measurement sessions and displays.

**Figure 1.**
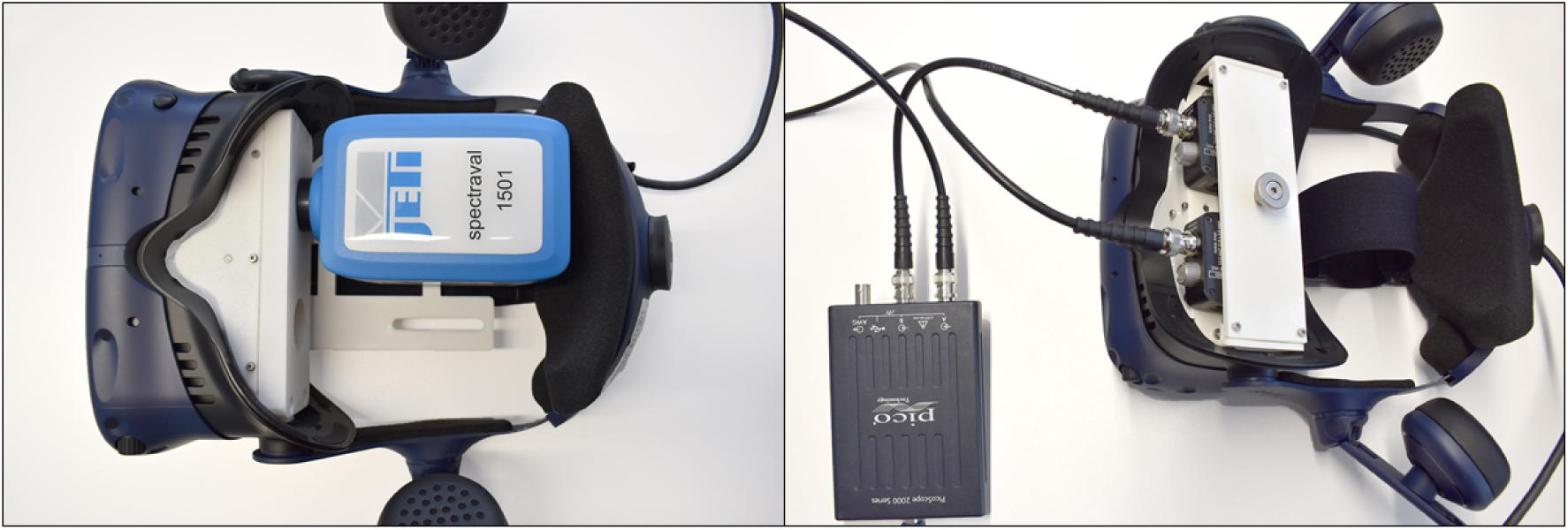
Photographs of the 3D-printed adapters used to make reproducible spectral and temporal calibration measurements across different HMDs.

Spectral irradiance data were collected for full-field white stimuli across several input levels (0 to 1 in 0.1 increments), with each measurement being repeated five times. The resulting spectra were used to derive the photopic illuminance (lux) and the alpha-opic equivalent daylight (D65) illuminance (lx) for the l-cone, m-cone, s-cone, rod and ipRGC photoreceptor cells.

Custom-written Python code was used to control the measurement process. This script handled gamma-corrected stimulus presentation in the headset, communicated with the spectroradiometer, and automated the acquisition of spectral data. Communication with the JETI spectroradiometer was implemented using the PyPlr toolbox (Martin et al., 2022), and illuminance and α-opic quantities were computed from the measured spectra using LuxPy (Smet, 2020).

#### Temporal calibration

A temporal calibration of the HTC Vive Pro Eye display was performed to ensure accurate delivery of the intended temporal light stimuli across a range of frequencies. Brightness measurements were recorded over time using a dual channel PicoScope oscilloscope (Pico Technology, St Neots, UK) connected to two mounted DET36A2 photodiodes from Thorlabs (Bergkirchen, Germany). The photodiodes were each placed in front of the respective eye or display of the VR headset, and were used to capture the light output and transform it into a voltage signal. The photodiodes were securely held in place using a custom 3D-printed adapter designed for the HTC Vive Pro Eye, ensuring precise and repeatable alignment with each eye display (see **Figure 1**, right). The design isolates the detectors and reduces stray light between lenses, supporting high-fidelity signal capture.

The measurements were taken at a sampling rate of 250 kHz for a duration of 20 seconds for each stimulus. The conditions tested included a white constant background and sinusoidal flicker at frequencies spaced logarithmically from 0.01 to 20 Hz (0.01, 0.02, 0.05, 0.1, 0.22, 0.5, 1, 2.24, 5, 10, and 20 Hz). A custom-written Python script was developed to generate and present these sinusoidal flicker frequencies simultaneously on both eyes of the display. The same script was also used to interface with the PicoScope oscilloscope, allowing automated signal acquisition and saving of the voltage responses. Communication with the oscilloscope was implemented using the picosdk Python library (Pico Technology Ltd., 2019).

The captured signals were analysed using the Fast Fourier transform implemented in Python to identify the frequency with the peak amplitude and compare it to the input flicker frequency. Additionally, the amplitude at harmonic frequencies was calculated to assess signal fidelity and distortion. The temporal modulation transfer function of the displays was also derived to characterize the system’s ability to reproduce different temporal frequencies accurately.

### Melatonin suppression study

Data for this study was collected as part of a larger study on binocular retinal signal integration in melatonin suppression (<u>ISRCTN88955347</u>). In addition to the two light conditions in this analysis, participants completed two additional experimental sessions with light conditions that varied temporally or spatially. The protocol followed was identical in all cases, except for five participants that received the light intervention with pharmacologically dilated pupils.

#### Participants

A total of 32 participants (age range 20-39 and mean ± SD: 27.2 ± 5.6; 12 males, 20 females) were included in this study. Participants were healthy, non-smoking individuals with no reported sleep disorders, substance abuse, medication or hormonal intake. All participants had normal or corrected-to-normal vision, confirmed through screening tests.

#### Procedure

Participants adhered to a standardized sleep-wake cycle, monitored using actigraphy (Condor ActTrust2, Condor Instruments, São Paolo, Brazil) and daily sleep diaries, for at least 7 days before their first laboratory session, and throughout the 3-weeks study. Alcohol and caffeine intake were restricted throughout the duration of the study. Participants visited the laboratory in four occasions, which were each at least 4 days apart, and the order of the stimuli presented in each session was randomized. After an evening session, participants were allowed to sleep one hour longer than usual to recover, and they returned to their habitual sleep schedule for subsequent days. When participants didn’t adhere to the regular sleep schedule before a session, the session was postponed or they were excluded from the study.

Participants arrived at the laboratory 5 hours before their habitual bedtime, underwent alcohol and THC screening, and completed the calibration procedure for the VR headset’s eye-tracking system, including adjustment of the headset position on the head, the distances between the two screens, and gaze tracking calibration.

Each experimental session began 4 hours before habitual bedtime. Participants sat in a dimly lit room (<10 lux) during the first 2 hours. Subsequently, they wore the VR headset to view either the dark or bright light stimulus for 20-minute intervals at a time in a 30-minute block, for four intervals in total. The remaining 10 minutes in each experimental block was used for saliva sample collection (2 minutes), a short auditory psychomotor vigilance test (3 minutes) and a 5-minute break. Experimenters were blinded to which light stimulus was being presented each session. While wearing the VR headset, participants were monitored using the integrated eye-tracker of the VR headset. An automated program monitored the openness of both eyes, and issued an audio warning if eyes were detected to be closed for more than 5 seconds. After 2 hours of light exposure through the VR headset, participants returned to the dimly lit room for 1 hour.

Saliva samples were collected every 30 minutes throughout the evening, directly after each light exposure block of 20 minutes. Following the saliva sample collection, participants took a break of 5 minutes. Some participants (n=5) viewed the stimuli with pharmacologically dilated pupils. For this, tropicamide eye drops were administered 30 minutes before the light intervention (Mydriaticum Stulln® UD 5 mg/1 ml, Pharma Stulln, Stulln, Germany).

During each evening session, up to 3 participants took part in the study at once. This was made possible by having three separate experimental bays separated by privacy screens, each equipped with an HTC Vive Pro Eye HMD. The use of VR headsets ensured that participants were not exposed to or influenced by the light stimuli presented to others, as each headset delivered individualized and controlled light stimuli directly to the participant’s eyes. This setup eliminated any potential cross-exposure to stray light, and ensured a constant light delivery throughout the intervention periods.

#### Stimuli

The light stimuli consisted of a constant background displayed uniformly across the field of view. In the dark condition, the intensity was set to zero, while in the bright light condition the intensity was set to 50% (after gamma correction and linearization of the display), corresponding to an average illuminance of 90.3 lux and melanopic EDI of 90.9 lx across both eyes of all headsets used. A fixation cross was positioned centrally to maintain participants’ gaze stability during light exposure.

#### Design

The study employed a within-subjects design, with each participant exposed to both dim and bright light conditions in separate sessions. Sessions were randomized and counterbalanced across participants. Data collection occurred in a controlled laboratory environment, ensuring consistent ambient conditions.

#### Data processing and analysis

Saliva samples (>1 mL) were collected using Salivettes (Sarstedt, Nürnberg, Germany) and were scanned and centrifuged immediately after collection. Samples were stored at -20°C and later analysed for melatonin concentration using enzyme-linked immunoabsorbent assay (ELISA) by NovoLytiX GmbH, Switzerland (MLTN-96, with typical specifications: limit of quantification 0.5 –50 pg/mL, detection limit 0.5 pg/mL, mean intra-assay precision: 7.8%, mean inter-assay precision 10.0%).

The melatonin data were processed in several steps using Python. Initially, sample order was verified based on the timestamps at the time of collection, and misordered samples were corrected (n=1 sample, total=704 samples). Missing melatonin concentrations were interpolated using the piecewise cubic hermite interpolating polynomial method, supplemented by forward and backward filling for remaining gaps (n=7 samples).

The area under the curve (AUC) for melatonin concentrations during light exposure were computed for each condition and participant using Simpson’s rule. Finally, percentage melatonin suppression was calculated relative to the dark condition for each participant as:

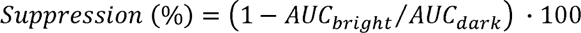

The statistical analysis was conducted using the lme4 package in R. We fitted a linear mixed-effects model to evaluate the effects of light condition, pupil dilation, and their interaction on normalized AUC values of melatonin concentrations. In addition to the fixed effects of condition and pupil dilation, a random intercept of participant was included to account for inter-individual variability in AUC values. The model incorporating the maximal random effects structure, including both random intercepts and slopes, was tested but failed to converge.

The model was fitted using restricted maximum likelihood estimation (REML). Parameter estimates and interaction effects were assessed, and 95% confidence intervals for the fixed effects were derived using a bootstrap method to provide robust interval estimates. Model diagnostics were performed to ensure validity, including residual analysis.

## Results

### Results of HMD calibration

We performed three types of measurements to characterize the display properties of the HTC Vive Pro Eye headset: luminance calibration and gamma correction, illuminance measurements, and temporal display profiling. For the illuminance and temporal measurements, we used custom 3D-printed adapters to ensure consistent and reproducible sensor positioning within the headset. This was particularly important for the illuminance measurements, which were performed across five different HTC Vive Pro Eye headsets, since this ensured the adapter maintained a fixed distance and angle between the spectroradiometer and the lens across all devices, enabling standardized and comparable data collection. To facilitate reuse and adaptation by other researchers, we have published the adapter designs under a permissive license that allows modification and redistribution, together with a technical note on their use (Fernandez-Alonso et al., 2025).

#### Luminance calibration and gamma correction

Luminance measurements were taken for both eyes of the HMD and for all individual primaries. As expected, we found a non-linear relationship between RGB input values and luminance output, which is standard in digital displays. We fitted the gamma curves to each individual RGB primary and to the white background to determine the correction factor. The results of the luminance measurements and the fitted curves are shown in **Figure 2**.

**Figure 2.**
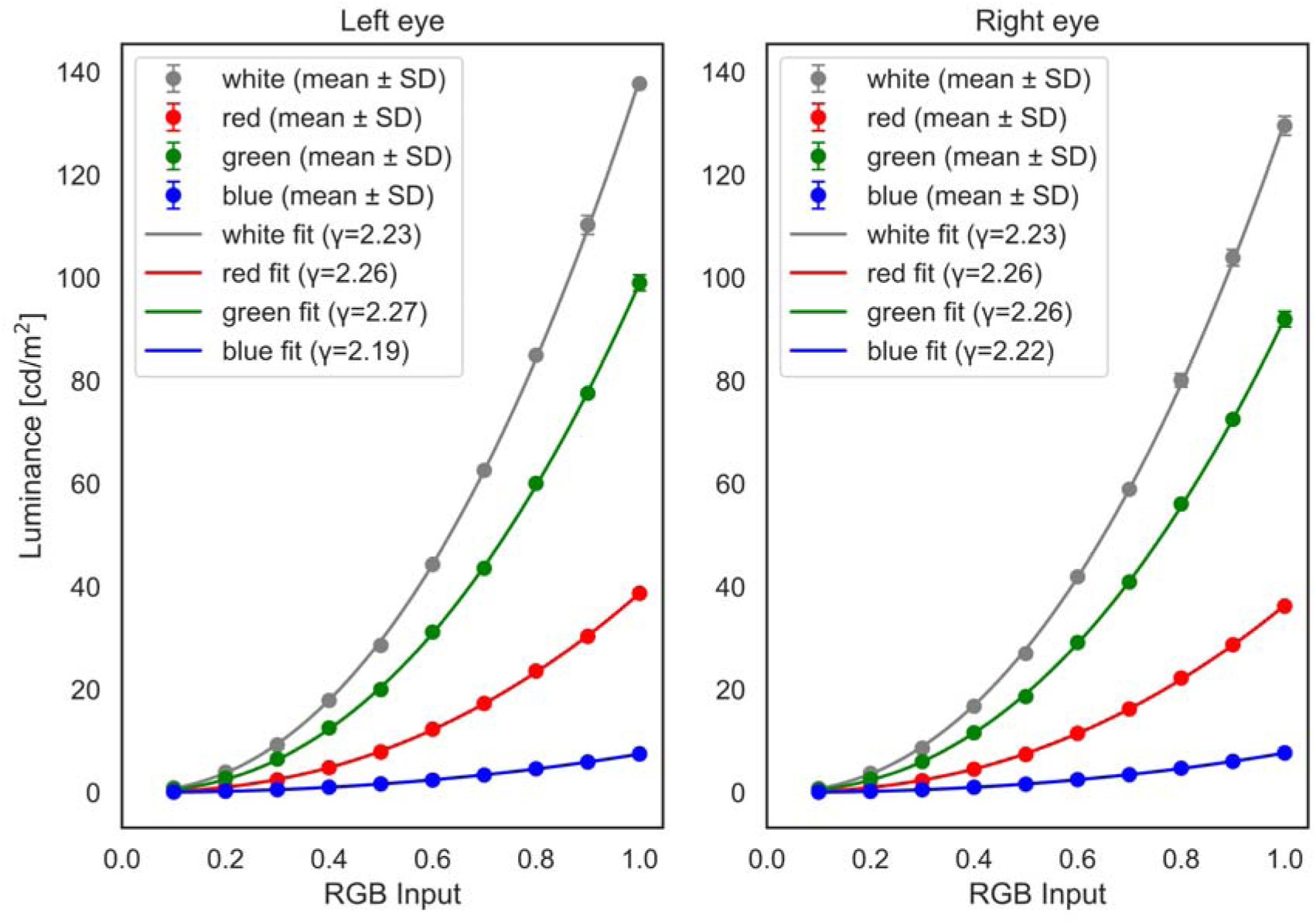
Luminance measurements and fitted gamma functions for each display primary and for white (equal RGB input). The round markers and error bars represent the mean and standard deviation of the measured luminance from three repetitions. The continuous lines represent the fitted power functions, with the estimated gamma values indicated in parenthesis on the legend.

As shown, the resulting gamma values are close to the standard used in most displays, 2.2. However, there are some discrepancies between primaries, particularly the blue colour has a shallower curve or lower gamma value than the red and green curves. While this difference is small, it could be relevant for studies where precise colour control of the stimulus is important.

Since for this study we used a white background as stimuli, we used the corresponding gamma value of 2.225, and calculated the correction factor as the inverse of this. After applying this correction, the luminance output was re-measured and compared against the target linear response. The results are shown in **Figure 3**. These luminance measurements were fitted with a linear regression, with a resulting coefficient of determination (R^2^) of 1, indicating that the gamma correction successfully linearized the luminance output of the display.

**Figure 3.**
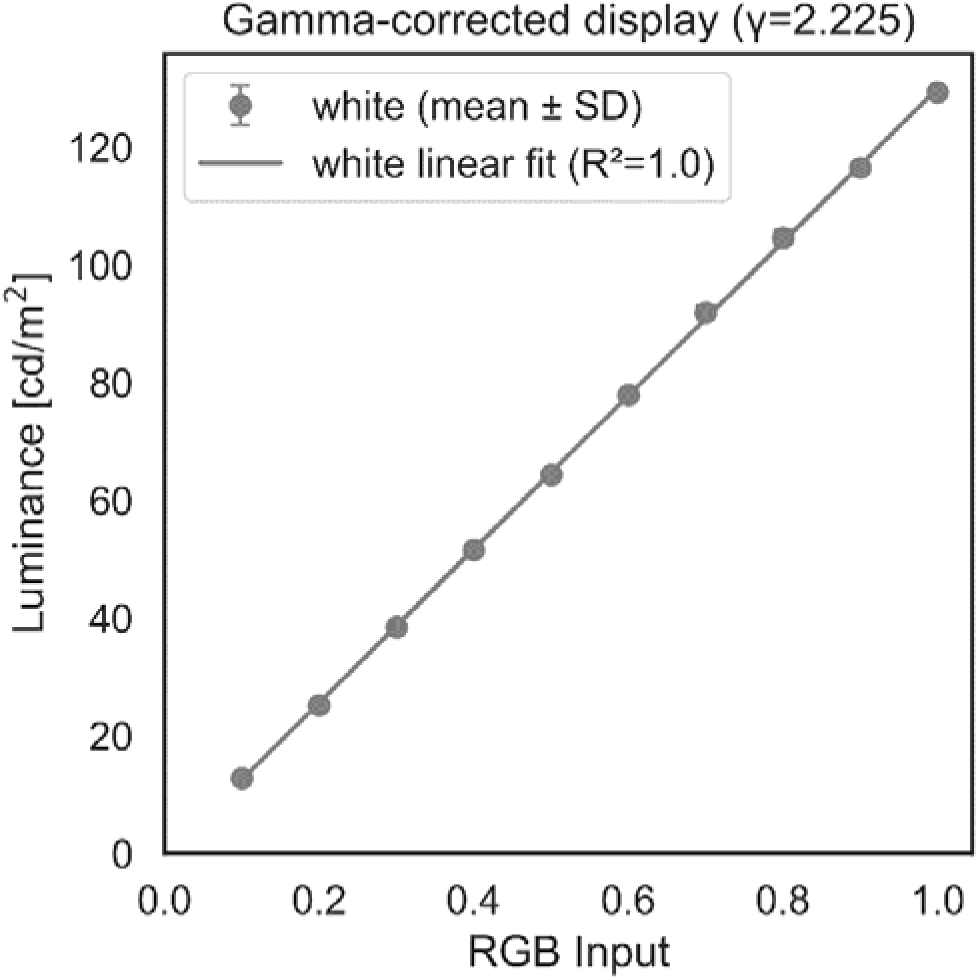
Measured luminance and fitted linear function after gamma correction of the display. The round markers and error bars represent the mean and standard deviation of the measured luminance from three repetitions. The continuous line represents the linear function fitted, with the R2 value of the fit shown in parenthesis on the legend.

#### Illuminance and α-opic equivalent measurements

We measured the spectral irradiance of the gamma-corrected HMDs across a range of input intensity levels (0 to 1 in steps of 0.1) and calculated the photopic and alpha-opic equivalent daylight (D65) illuminance (lx) for the l-cone, m-cone, s-cone, rod and ipRGC photoreceptors. Measurements were conducted on five different HTC Vive Pro Eye headsets to evaluate inter-device consistency. The use of a 3D-printed adapter ensured that the spectroradiometer was positioned at a fixed distance and angle relative to the lens across all headsets, enabling standardized measurements.

Figure 4 shows the mean photopic illuminance and mean α-opic equivalent daylight illuminance (EDI) values (melanopic, rhodopic, L-cone-opic, M-cone-opic, and S-cone-opic) plotted against input intensity for the left and right eyes of the HMDs. These results are also available in Supplementary Table 1. As expected from gamma-corrected displays, all metrics increase linearly with input level. The relative proportions of the α-opic channels remained consistent across headsets, indicating spectral stability of the luminous output. The variability among the different headsets is low, except for a lower illuminance in the right eye of headsets 3 and 5. Due to the standardized adapter used, this is unlikely to be solely due to the measurement setup, and more likely reflects real variability in the optical output or transmission characteristics of the right-eye display panels or lenses in those specific headsets.

**Figure 4.**
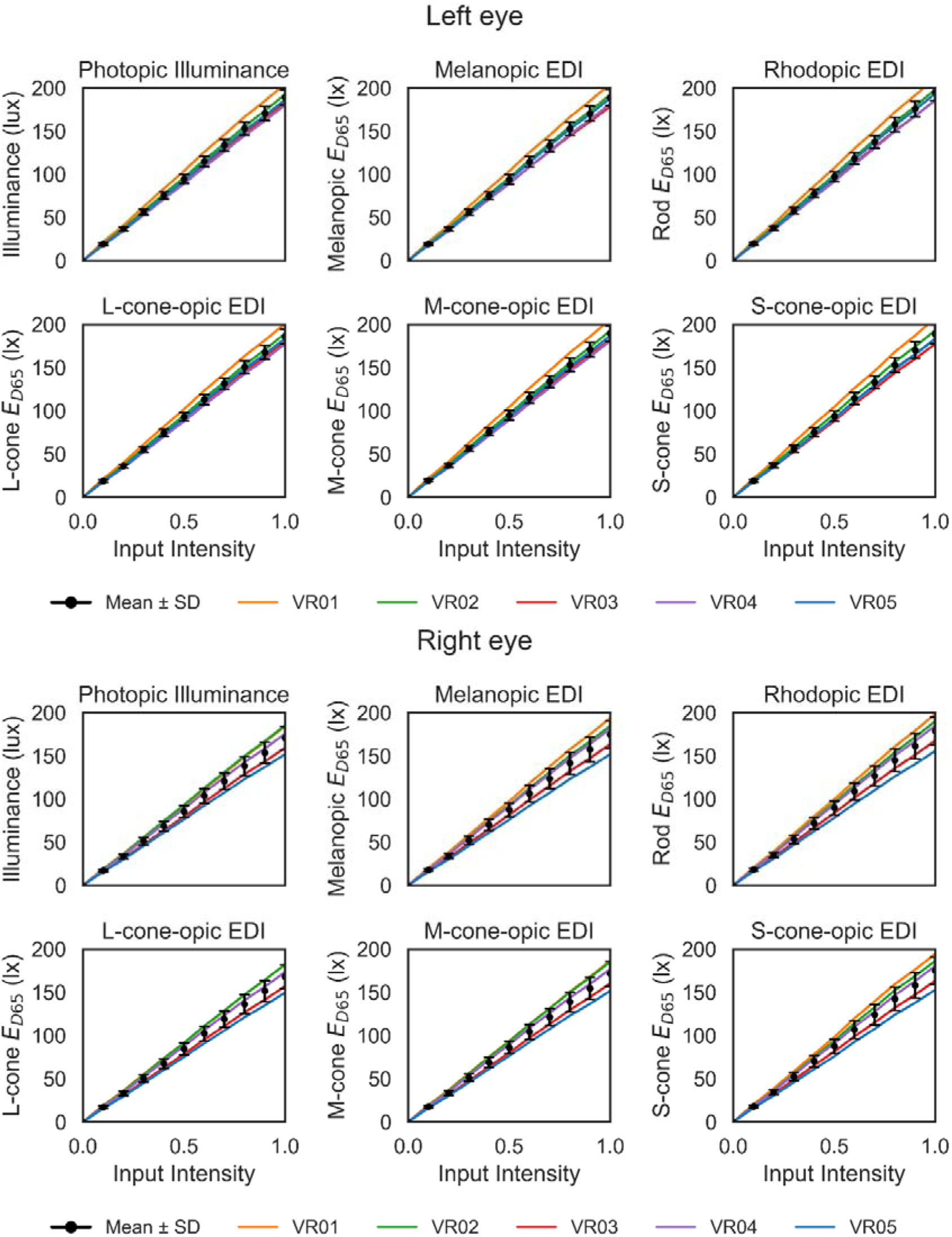
Photopic and α-opic equivalent daylight (D65) illuminance for the ipRGC, rod, l-cone, m-cone, and s-cone photoreceptors as a function of input intensity, plotted separately for the left (top) and right (bottom) eyes of the HTC Vive Pro Eye HMDs. Individual lines represent data from five different headsets as indicated by the legend, while the back circles and error bars indicate the mean ± standard deviation (SD) across all headsets for each eye.

#### Temporal calibration

The luminance of the display was measured for both eyes with high temporal resolution, while displaying sinusoidal flicker at a range of temporal frequencies (from 0.01 to 20 Hz) and for a constant white background. For the latter, the measurements revealed that the display has a constant nearly sinusoidal strobing of the emitted light at 90 Hz, illustrated in Figure 5. According to information from the manufacturer, this feature is encoded in the driver of the display and cannot be disabled. Therefore, a peak amplitude at 90 Hz was found for all other measurements taken.

**Figure 5.**
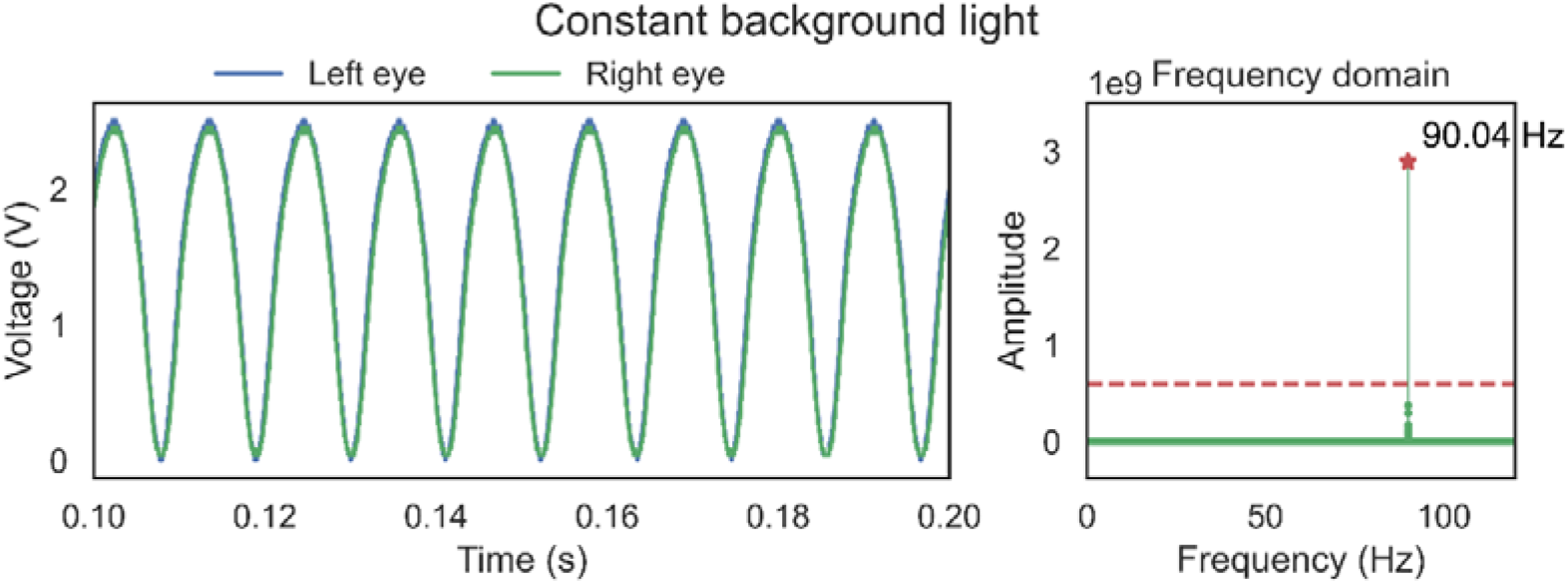
Measured signal in the time domain (left) and in the frequency domain (right) for a constant white background in the left eye and right eye of the display (note that both lines overlap significantly). The dashed red line on the right panel indicates the threshold used to determine the peak amplitude. The frequency with peak amplitude is shown as an annotation (90.04 Hz)

In Figure 6 we illustrate the results of measurements taken for sinusoidal flicker of 5 Hz. As seen, the strobing of the display at 90 Hz is still present, but the target frequency is displayed with high accuracy and low distortion, as indicated by low power at harmonic frequencies.

**Figure 6.**
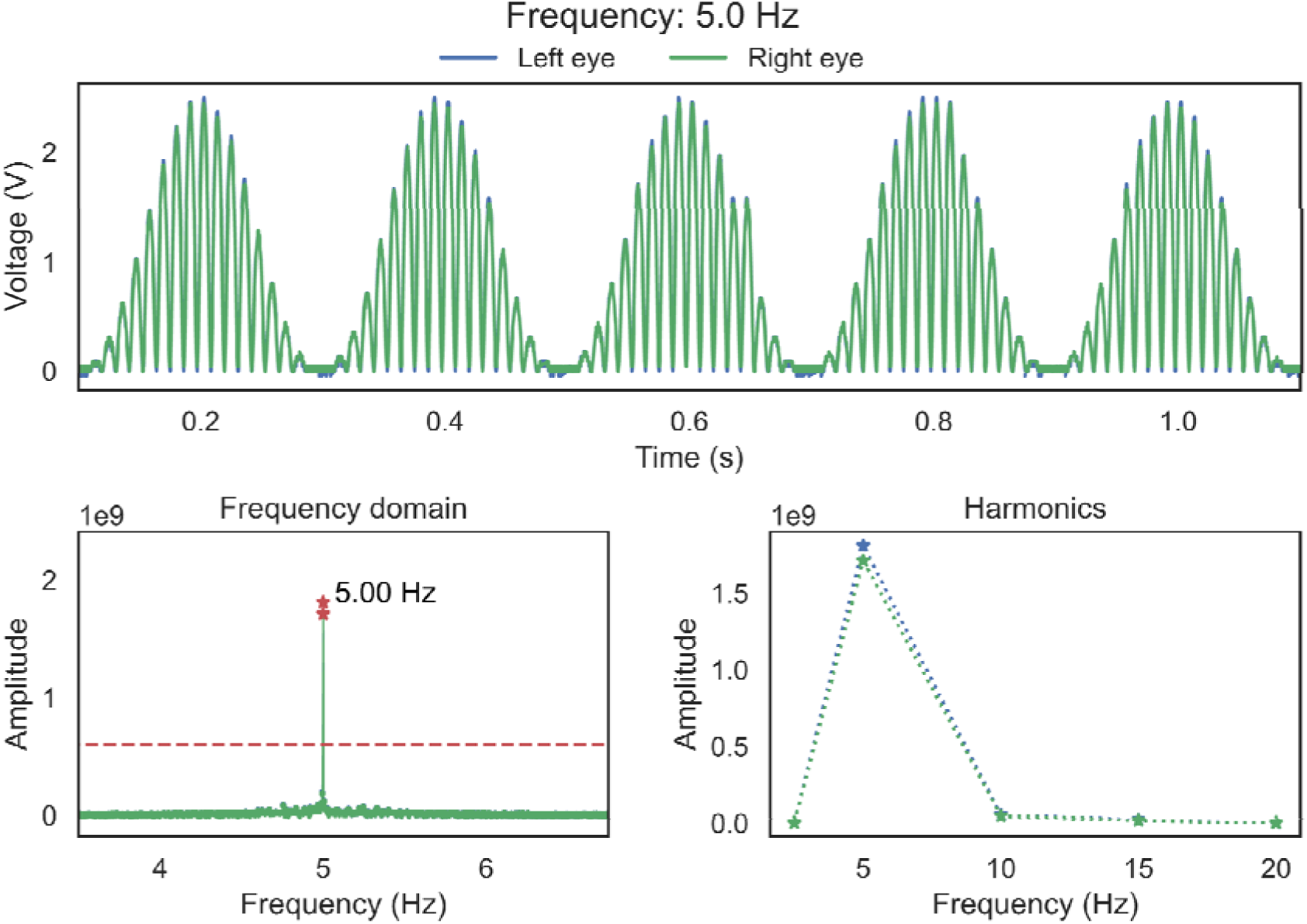
Measured signal for sinusoidal flicker of 5 Hz shown in the left eye and right eye of the display in the time domain (top) and the frequency domain (bottom left). The amplitude at the harmonics of the peak amplitude frequency is shown (bottom right). The dashed red line on the bottom left panel indicates the threshold used to determine the peak amplitude. The frequency with peak amplitude is shown as an annotation on the bottom left figure. Note that the data of both eyes (blue and green) overlap significantly,

Figure 7 summarizes the results of all frequencies tested, displaying temporal transfer function and the amplitude at the harmonics of the target frequencies. Despite the inbuilt strobing, the display was able to reproduce sinusoidal flicker with high accuracy, minimal distortion, and high agreement between both eyes across the measured frequencies, although a significant decrease in amplitude was observed at 20 Hz, reducing the fidelity of the displayed stimuli.

**Figure 7.**
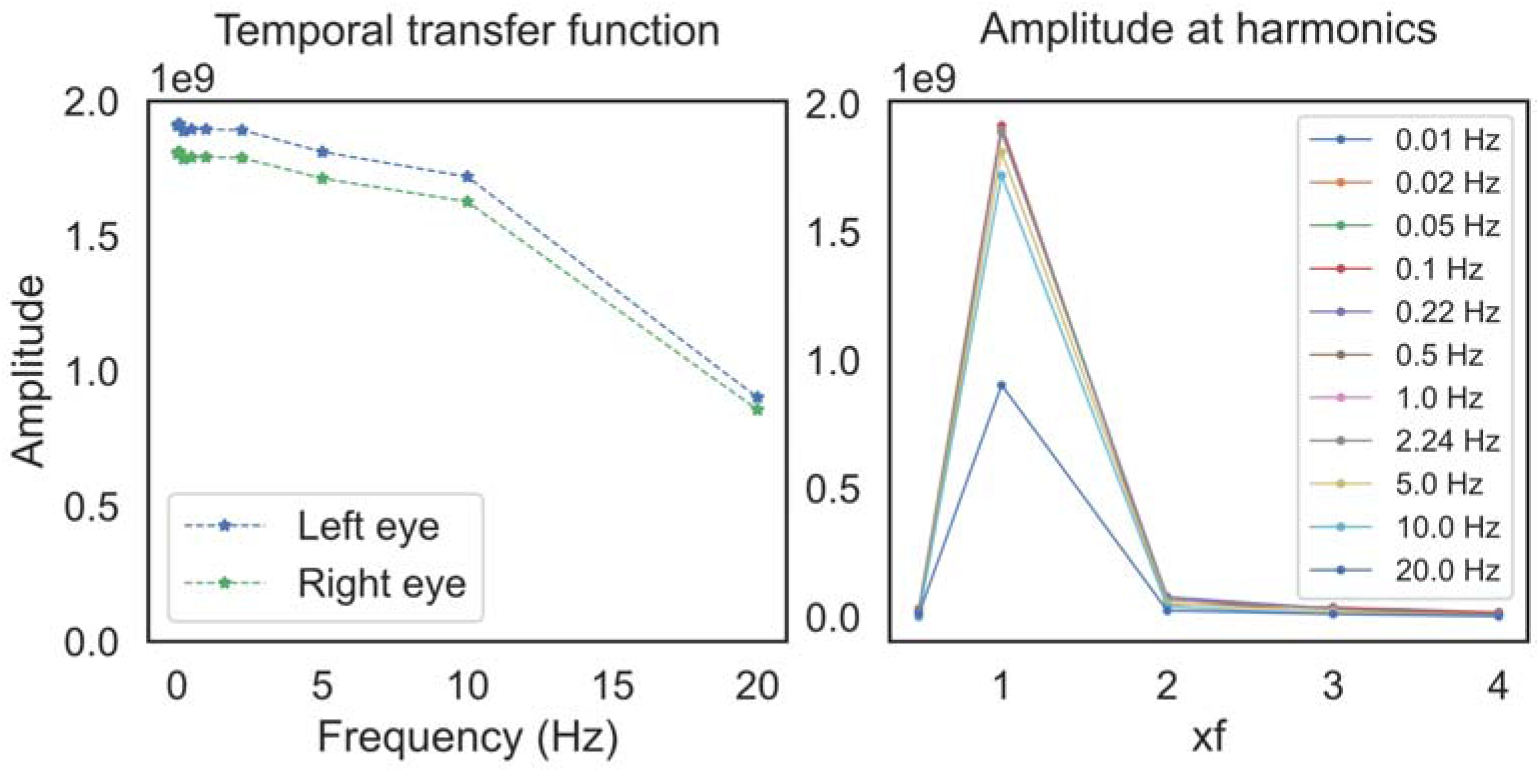
Temporal modulation transfer function of the display (left) and amplitude at the harmonics of each measured frequency.

We also tested higher frequencies such as 25 Hz, but there the interaction between the sinusoidal flicker and the inbuilt strobing severely distorted the waveform, causing high amplitude at higher harmonics. It is possible that this issue could be ameliorated by using a different waveform such as square-wave flicker; although our findings in general indicate that the display is best suited for delivering temporally-modulated stimuli at low to mid-range temporal frequencies.

### Results of melatonin suppression study

We investigated the acute melatonin suppression effects of light stimuli delivered with head-mounted virtual-reality displays in 32 healthy adults. We tested two conditions: a dark background, and a constant bright light background of 50% intensity, which corresponds to an average illuminance of 90.3 lux and average melanopic EDI of 90.9 lx across both eyes and all headsets used. An example of the results for one participant are shown in Figure 8.

**Figure 8.**
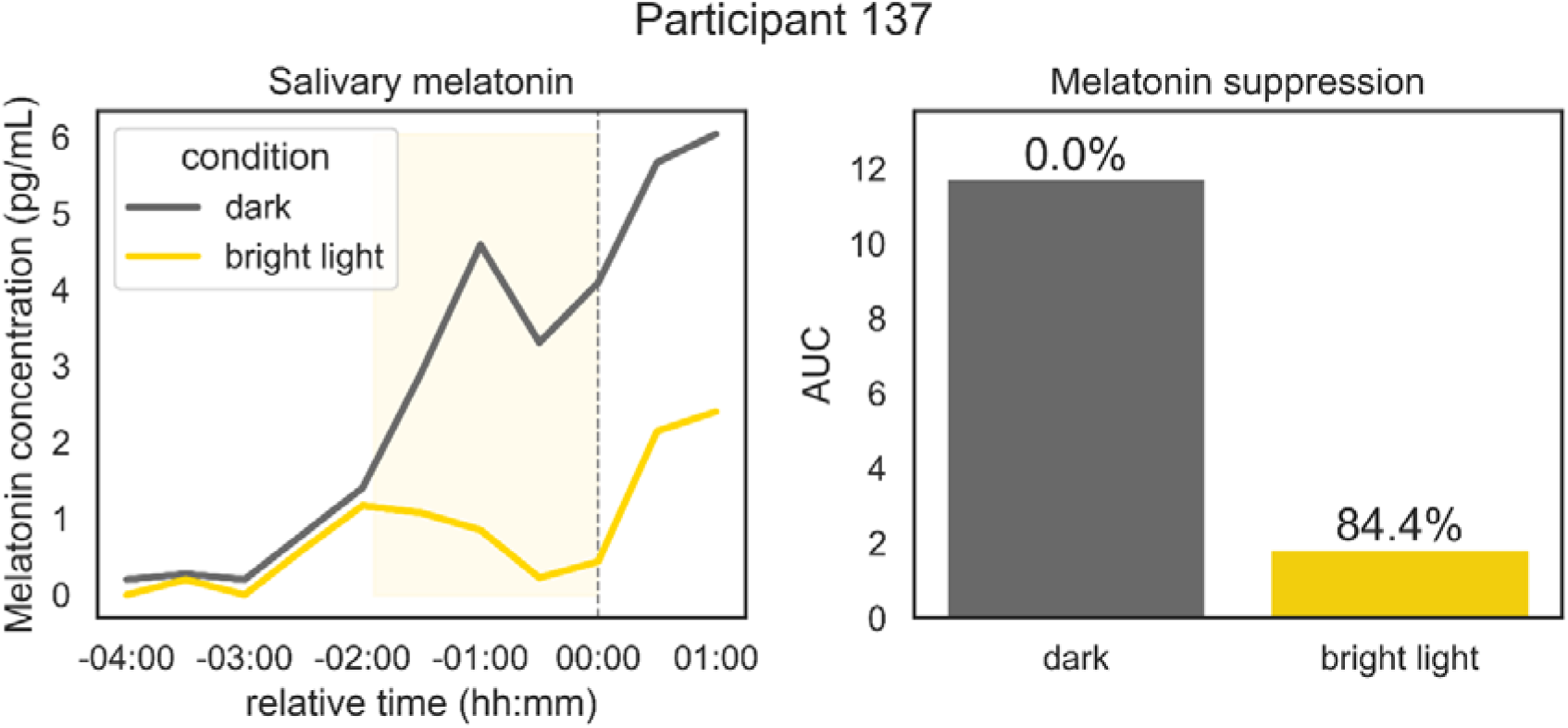
Salivary melatonin concentration as a function of time relative to habitual bedtime (left) and the calculated AUCs during light exposure for each condition (right) for participant 137. The labels above the bar plots represent the percent of melatonin suppression calculated as a function of the dark condition.

Figure 9 shows the results for all participants in the sample, separated by whether participant’s pupils were pharmacologically dilated or not. As shown on the left panel of Figure 9, the cumulative melatonin concentration during light exposure is lower in the bright light condition for 24 out of 32 participants (75%) compared to the dark condition, indicating an acute suppression of melatonin.

**Figure 9.**
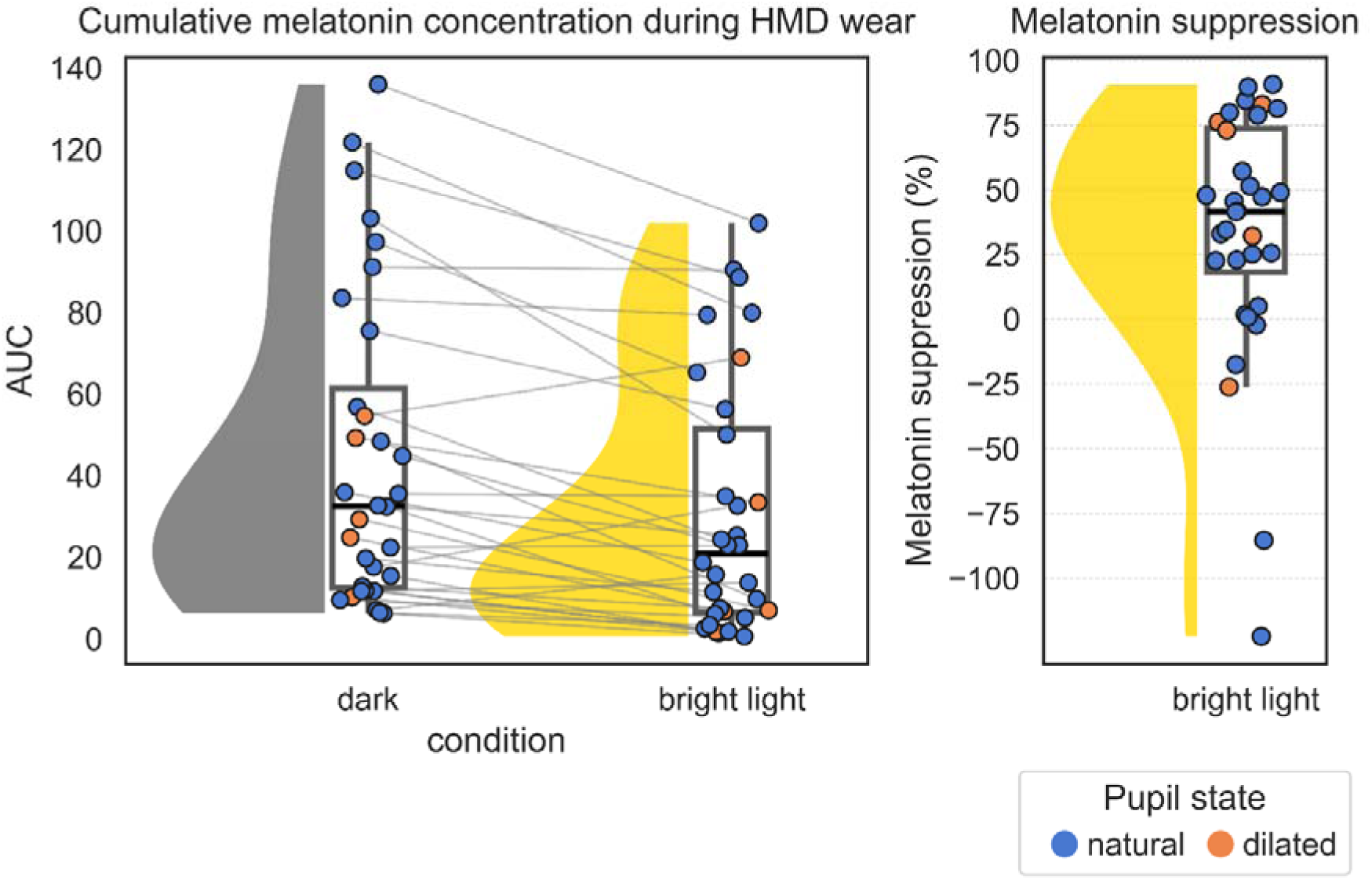
Cumulative melatonin concentration as a function of condition (left) and calculated melatonin suppression for the bright light condition (right). Each marker represents one individual participant. The colour of the markers indicates if the light stimulation was received with a natural or pharmacologically dilated pupil. The continuous grey lines indicate repeated measures, connecting the data points that belong to the same participant. The smoothed filled curves represent the estimated distribution of the data obtained through kernel density estimation. The black box plot indicates the median, the 25th (first quartile), and the 75th (third quartile) percentiles, with whiskers extending to represent approximately 1.5 times the interquartile range.

To examine this effect statistically, we fitted a linear mixed model on the AUCs, with a fixed effect of condition, pupil dilation, and their interaction, and a random intercept of participant. This means that the baseline AUC in the dark condition was allowed to vary randomly for each participant. A model with the maximal random effects structure (i.e., with a random slope of condition) was also tested, but failed to converge. The results of the model fit are shown in **Table 1**. As seen, there is a significant effect of experimental condition, with the estimated AUC under the bright light condition being significantly lower than under the dark condition by 14.41 (CI 95% -20.76 to -8.76, *p < 0.001*). Regarding pupil dilation, no significant effects were found, neither in the dark condition nor in the bright light condition. This means that participants that took part in the experiment with pupil dilation did not differ significantly from the participants with natural pupils, and that being exposed to the bright light condition did not have any significant further effects in the AUC.

**Table 1.**
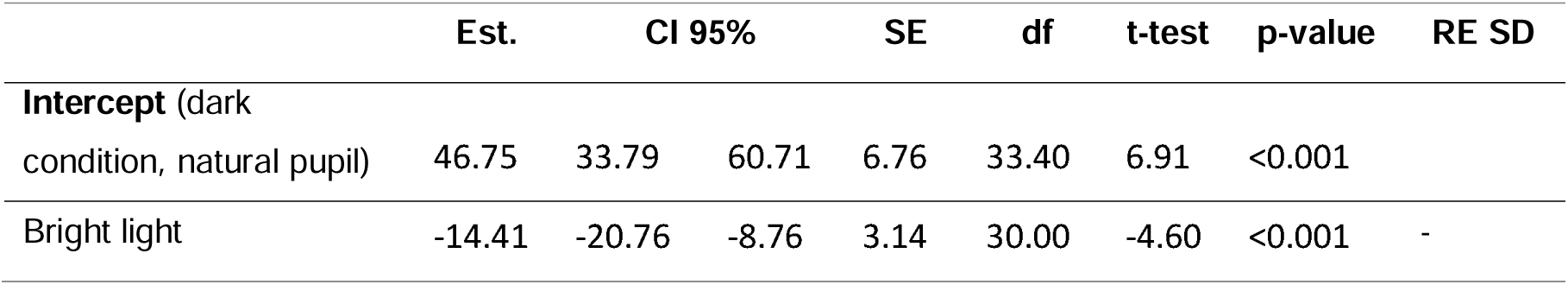
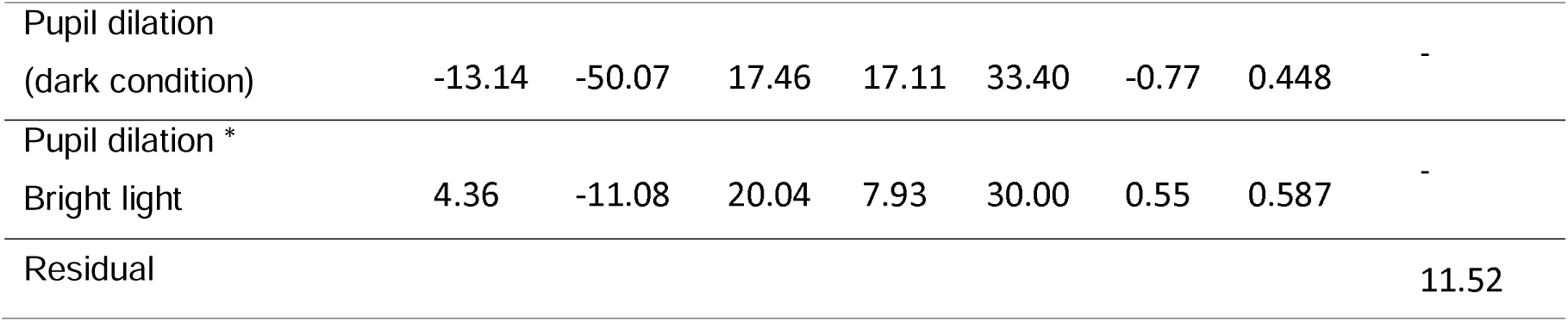
Results of the linear mixed model fit of cumulative melatonin concentration (AUC) during light exposure with fixed effects of condition, pupil state and their interaction, and random intercept of participant. The results shown are the parameter estimates (Est.), the 95% confidence intervals (CI 95%), the standard error (SE), degrees of freedom (df), t-test and p-values, as well as the standard deviation of the random effects (RE SD)

To better illustrate what this represents in terms of melatonin suppression, we show in the right panel of Figure 9 the distribution of the suppression percent in the sample, and we provide descriptive statistics in **Table 2**. On average, melatonin was suppressed by 33.3% in the bright light condition for all participants. When comparing the subgroups by pupil state, we see that those with dilated pupils had a higher mean melatonin suppression at 47.6%, compared to 30.7 % in the natural pupil group. However, in both groups we see considerable variability in the melatonin suppression effect with some extreme outlier values, which could explain the lack of statistically significant differences between both groups in the previous analysis.

**Table 2.** Descriptive statistics of melatonin suppression percent for participants that received the light stimuli with dilated or natural pupils, and for all participants.

|  | Mean | SD | Min | 25% | Median | 75% | Max |
| --- | --- | --- | --- | --- | --- | --- | --- |
| **Dilated pupil** | 47.55% | 45.83% | -26.28% | 32.07% | 73.00% | 76.08% | 82.90% |
| **Natural pupil** | 30.69% | 49.10% | -122.38% | 13.73% | 41.42% | 54.31% | 90.61% |
| **All participants** | 33.33% | 48.28% | -122.38% | 18.12% | 41.58% | 73.77% | 90.61% |

#### Pupillometry results

Pupil diameter was extracted from the inbuilt eye-tracker of the HMD to assess light-induced pupil responses. The results are show in Figure 10. As expected, pupil diameter was larger in the dark condition compared to the bright light condition, consistent with normal pupillary constriction to light. Participants who had pharmacologically dilated pupils showed pupil sizes comparable to those in the dark condition.

**Figure 10.**
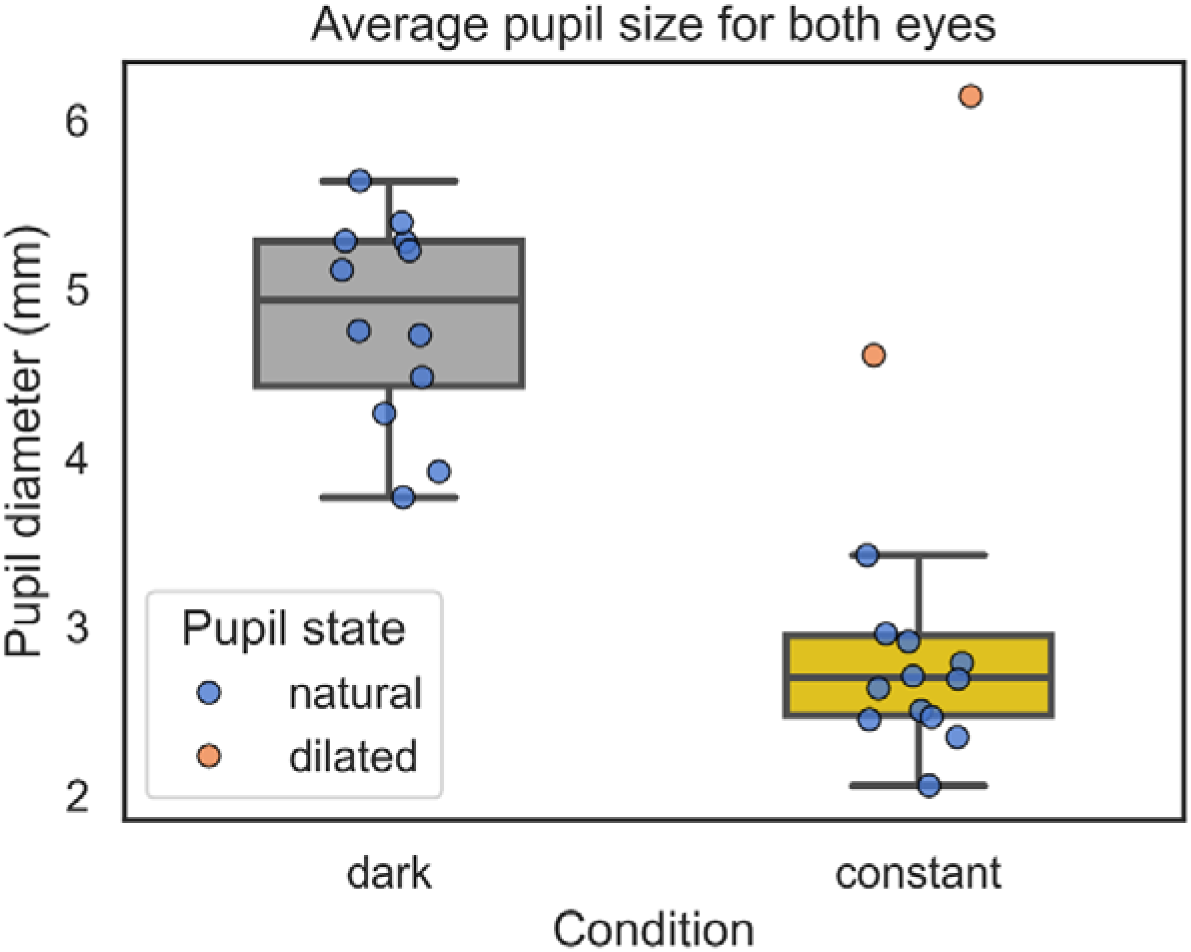
Mean pupil diameter across both eyes for each participant during both light conditions. Boxplots represent the group-level distribution, while individual participants are shown as circles coloured by pupil state.

Notably, the eye-tracking system failed to provide usable data in 34 sessions out of 60 sessions where eye-tracking was used (57%). In our experiment, the eye-tracker was only calibrated once at the beginning of the session, after which participants took the headset off. This could indicate that optimal tracking performance requires recalibration each time the headset is worn.

## Discussion

In the present study, we demonstrated that head-mounted virtual-reality displays can be effectively calibrated to deliver controlled light stimuli for chronobiology research. First, we established luminance calibration and gamma correction procedures, verifying that the non-linear relationship between digital input values and luminance output could be successfully linearized. Despite minor discrepancies between primary colours—particularly for blue—our final gamma correction achieved a near-perfect linear response (R^2^ = 1). Second, temporal calibration revealed the presence of a built-in 90 Hz strobing in the display driver, which was not user-accessible for deactivation. Nonetheless, the system reproduced sinusoidal flicker at low-to-mid temporal frequencies (up to 10 Hz) with minimal distortion.

We also completed a melatonin suppression study involving 32 healthy adults, and we observed a significant reduction in melatonin levels under a bright light condition (white background of approximately 90.3 lux illuminance and 90.6 lx melanopic EDI) compared to a dark condition (black background of approximately 0.0 lux illuminance and 0.0 lx melanopic EDI). This suppression reached an average of 33.3% across participants, demonstrating that VR displays can elicit robust acute non-visual light responses. Increasing the brightness of the screen to 100% would be expected to produce an even greater suppression effect (Prayag et al., 2019). No participants raised any concerns about the comfort in wearing the headsets, indicating tolerability of the stimulation method.

Contrary to previous findings (Gaddy et al., 1993), the use of pharmacological pupil dilation did not result in a statistically significant difference in melatonin suppression, likely due to substantial inter-individual variability and the small number of subjects that underwent pupil dilation.

### Advantages of head mounted displays

#### Enhanced precision and reduced variability

The calibration procedures demonstrated that VR HMDs can provide high-precision control over both the spectral and temporal properties of the light stimulus. This level of consistency surpasses what is usually achievable with traditional setups, where factors such as viewing angle and room reflections can introduce unwanted variability in retinal illumination (Spitschan, 2021). By delivering light directly through an HMD, each participant receives the same stimulus, regardless of posture or external conditions.

#### Automation and eye-tracking integration

An additional benefit of this system lies in the integration of real-time eye tracking. By monitoring participants with an automated program, the experimenters were able to confirm that participants were indeed viewing the stimulus, thereby reducing the need for constant supervision. In general, the programmability of HMDs supported a high degree of automation in this study. We employed custom software to deliver audio instructions throughout the experiment and precisely schedule steps like light exposure and saliva sampling. The use of audio instructions and the integration of an external response box, allowed participants to advance through the experimental protocol on their own by pressing a button, giving them more autonomy and removing the need for continuous step-by-step instructions from the experimenter. This high degree of automation significantly reduced the burden on experimenters and minimized errors from manual intervention. Another advantage was the possibility to test multiple participants in parallel without any risk of cross-exposure to stray light from other participants taking part. All these factors vastly accelerated the entire data-gathering process.

#### Experimental design and stimulus variety

The programmability of the displays also allowed us to present more complex stimuli as part of a larger study, such as temporally modulated light with different inputs to each eye. Other possibilities would include presenting spatially modulated stimuli, or even performing more complex visual tasks such as visual attention tests. Another added benefit was the possibility to randomize the order of the conditions while experimenters remained blinded to which stimulus was being presented each session.

#### Participant comfort

The use of HMDs also gave participants greater freedom of movement while being exposed to the stimulus, increasing comfort. Traditional highly controlled lab-based setups such as the ganzfeld dome stimulator, often require participants to remain in a fixed position in front of a light source throughout the duration of the intervention. This constraint is not present HMDs, allowing participants to sit more comfortably and to change their posture, while maintaining the same visual stimulation. This is particularly key for circadian experiments due to the long duration of experimental protocols.

### Implications for circadian research

Our findings confirm that VR HMD-based light delivery can effectively induce acute melatonin suppression, a key endpoint in circadian and sleep research. By precisely regulating light intensity and spectral composition, researchers can better isolate the physiological effects of specific lighting conditions. This method also opens the door to investigating advanced protocols that incorporate flicker or other temporally or spatially varying light stimuli, potentially providing insight into how dynamic lighting patterns influence circadian phase and other physiological rhythms.

#### Standardized stimuli across multi-centric studies and projects

The proposed approach facilitates the standardization of stimuli across different study sites. It is generally non-trivial to equate stimulus conditions across different laboratory settings and technological specifications. For example, when using room illumination as a stimulus, the exact dimensions of a room and light-related parameters such as the wall reflectance will modify the corneal light exposure received by a participant. Even when using integrating spheres with light sources, there is typically no guarantee that the stimuli received by an observer will be the same across two different setups. Using standard consumer hardware to display stimuli solves this issue, thereby opening the door to collect harmonized data linked by a common stimulus display system across multiple sites.

#### Higher efficiency in in-laboratory data collection

A key feature of experiments targeting neuroendocrine and circadian physiology is that they are costly in terms of time and resources. Using VR displays makes the collection of data a lot more efficient, due to the possibilities for automation and the ability to run multiple participants at a time.

#### Scalability for remote and at-home data collection

An especially promising avenue is the use of VR HMDs for data collection outside the laboratory. When combined with user-friendly calibration protocols and remote monitoring (e.g., via embedded eye tracking or integrated sensors), participants could complete lighting studies and interventions at home with scientifically validated stimuli. This approach would facilitate large-scale studies and longitudinal designs, expanding research capacity to populations that may otherwise be unable or unwilling to travel to specialized research facilities.

### Limitations and future directions

Although the present study demonstrates the feasibility of using VR displays for controlled light delivery, certain limitations must be addressed. First, the inbuilt strobing at 90 Hz found in the devices used, which could not be disabled, poses challenges for high-frequency flicker stimuli above approximately 10 Hz. Other headset models might not have this feature or offer options for disabling it for improved temporal precision.

We also found minor discrepancies in the gamma function of the different colour primaries, which could mean that using these displays is less suitable for studies where high accuracy of the individual display primaries is needed, such as for the silent substitution method. However, this could be addressed with more extensive light measurements and a more complex calibration procedure.

There was a subset of participants for which the light stimulation did not produce an acute suppression effect on melatonin. This could be addressed by using a higher intensity, since the stimulus used in this study was only 50% of the maximum brightness of the HMD. Potentially, the use of pupil dilation could also increase the suppression effect by increasing retinal illuminance (Gaddy et al., 1993), although this increase was not statistically significant within our sample.

Beyond these limitations found in our study, other potential limitations of HMDs require careful consideration. While the HMD used in this study allowed for individual adjustment of the interpupillary distance, and this was a required step as part of the inbuilt calibration procedure, this feature is not present in all commercial devices. This could introduce discrepancies in the visual stimulus received by different participants. In general, establishing standardized fitting protocols and using robust calibration procedures, together with providing clear adjustment instructions to participants, are necessary steps in order to ensure the precision of this methodology. Another characteristic of most HMDs is that they require straps to be secure around participant’s heads, making them rather unsuitable for studies that use electroencephalogram, facial electromyography, or other similar techniques that require free access to the face or the head.

Using the proposed HMD-based stimulation method, stimuli can be closely controlled for mechanistic studies, representing an effective stimulus to the ipRGCs and downstream neuroendocrine pathways. However, stimuli delivered in HMDs can differ markedly in the extent to which they represent the real world. Presenting naturalistic stimuli while carefully controlling it parametrically is an attractive outlook for future research.

Finally, future research should assess a larger variety of commercially available devices. Different VR headsets have varying hardware specifications and different level of control over the image being presented. Furthermore, different models would fundamentally differ in their level of brightness, resolution and field of view. Future work should assess which devices among those available are suitable for use in research, particularly for multi-site or large-scale studies.

## Conclusion

Our results demonstrate the promising potential of commercial head-mounted displays as a highly controlled, scalable, and standardized tool for investigating non-visual effects of light such as melatonin suppression. By combining meticulous temporal and spectral calibration and the capacity for multi-site and remote data collection, VR displays can significantly enhance the methodological capabilities of chronobiology and sleep research.

## Supplementary Materials

**Table S1.**
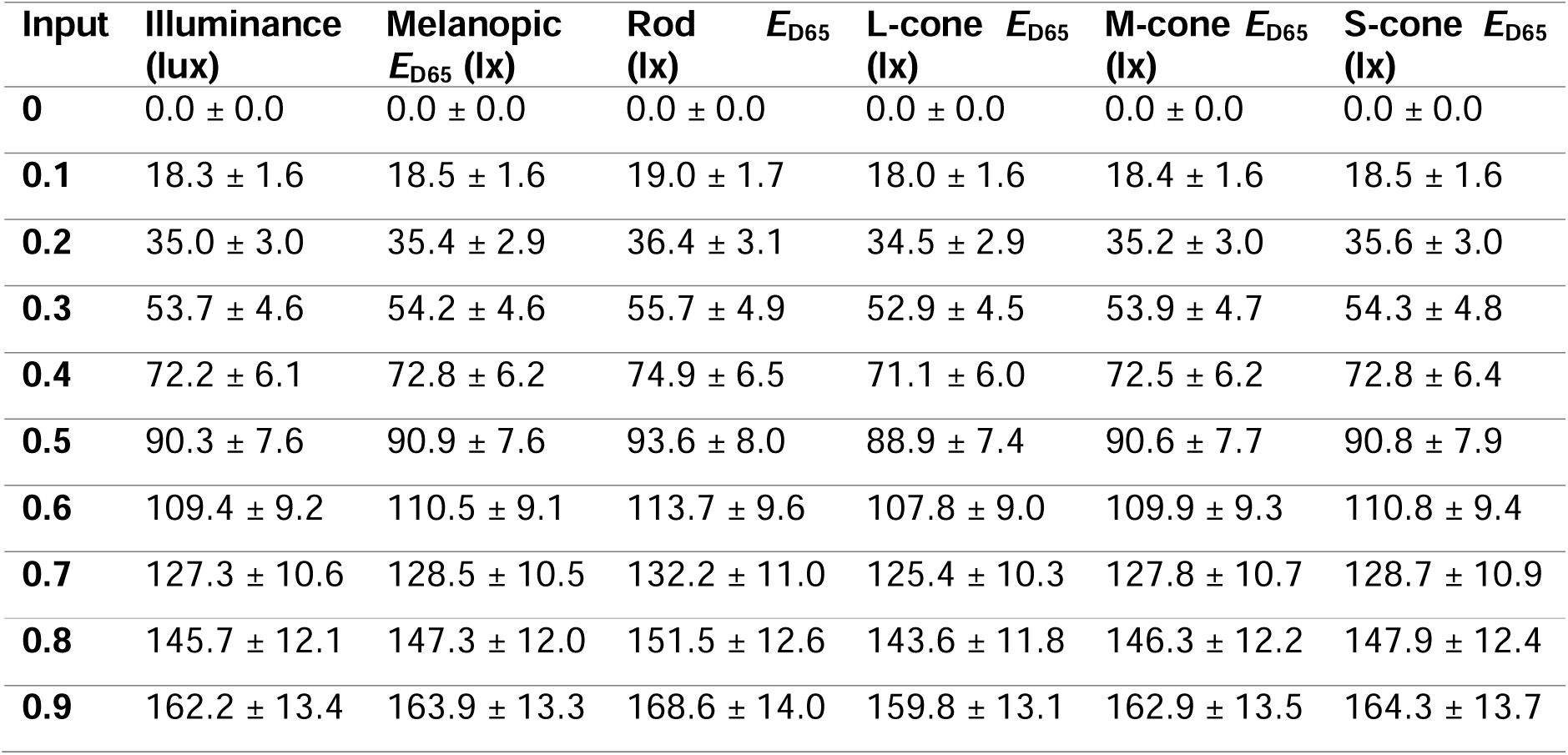
Mean ± standard deviation of photopic and α-opic equivalent daylight (D65) illuminance values across five VR headsets, averaged over both eyes, for each input intensity level.

## Author contributions

Conceptualization, M.F.A. and M.S.; Methodology, M.F.A. and M.S.; Software, M.F.A.; Validation, M.F.A. and M.S.; Formal Analysis, M.F.A.; Investigation, M.F.A. and M.S.; Resources, M.S.; Data Curation, M.F.A; Writing – Original Draft Preparation, M.F.A.; Writing – Review & Editing, M.F.A. and M.S.; Visualization, M.F.A.; Supervision, M.S.; Project Administration, M.F.A. and M.S.; Funding Acquisition, M.S.

## Ethics statement

The study was conducted according to the guidelines of the Declaration of Helsinki, and approved by the Ethics Committee of the Technical University of Munich (2022-439-S-SR).

## Acknowledgments

We thank Jennifer Reif, Nik Novik and Sing Yik Chan for their contributions to this project. Furthermore, we thank the staff at the mechanical and electronics workshop of the Max Planck Institute for Biological Cybernetics for their technical support.

